# Long-term repeatability in social behaviours suggests stable social phenotypes in wild chimpanzees

**DOI:** 10.1101/2020.02.12.945857

**Authors:** Patrick J. Tkaczynski, Alexander Mielke, Liran Samuni, Anna Preis, Roman Wittig, Catherine Crockford

## Abstract

Animals living in social groups navigate challenges when competing and cooperating with other group members. Changes in demographics, dominance hierarchies or ecological factors, such as food availability or disease prevalence, are expected to influence decision-making processes regarding social interactions. Therefore, it could be expected individuals show flexibility in social behaviour over time to maximise the fitness benefits of social living. To date, research across species has shown that stable inter-individual differences in social behaviour exist, but mostly over relatively short data collection time periods. Using data spanning over 20 years, we demonstrate that multiple social behaviours are repeatable over the long-term in wild chimpanzees, a long-lived species occupying a complex fission-fusion society. We controlled for temporal, ecological and demographic changes, limiting pseudo-repeatability. We conclude that chimpanzees living in natural ecological settings have relatively stable long-term social phenotypes over years that may be independent of life history stage or strategies. Our results add to the growing body of literature suggesting consistent individual differences in social tendencies are more likely the rule rather than the exception in group-living animals.

## Introduction

The fitness benefits of social bonds and social connectivity are well established across group-living animals, including in humans (1–11). Despite the adaptive advantages of maintaining social relationships, short-term strategies of social interaction avoidance could also improve fitness, such as during periods of disease outbreak (12). Similarly, during periods of social upheaval, such as instability in dominance hierarchies, individuals may change how they distribute their affiliative or aggressive social investment to reinforce key relationships or dominance position respectively (13,14). Fluctuations in resource availability (e.g. number of available mating partners or food availability) may also influence time allocation to affiliative social interactions, rates of aggression, social partner choice or general gregariousness (15,16). Lastly, an individual’s physiological state may fluctuate over time, e.g. during pregnancy and/or the rearing of offspring, in turn influencing motivation for social behaviours (17).

Substantial variation in socioecological settings and internal state suggests individuals should show flexibility in their social interaction patterns (18). However, consistent individual differences in social behaviour have been identified across various animal taxa, suggesting group-living individuals tend to show stable tendencies in solving social problems and interacting with other group members (19–26). This raises the question of why some individuals appear to be consistently more or less cooperative, gregarious, or aggressive than others.

A high degree of repeatability in a trait, i.e. the proportion of variation attributable to between-individual differences (27), may reflect underlying stable factors such as genetics and/or irreversible developmental adjustment to early life conditions (28–32). Under such a framework, individual patterns of social behaviour, and thus social phenotypes, emerge as a consequence of the interaction between genetics and exposure to the physical and social environment during development (30,32). The “social niche hypothesis” specifically proposes that consistent individual differences in behaviour arise due to niche specialisation to ameliorate within-species and/or within-group competition for resources (33). Here, an individual adopts a behavioural or social strategy to acquire resources dependent upon its existing competition-related characteristics, such as body size, health, dominance etc.

Alternatively, consistent between-individual differences in behaviour and sociality may arise due to behavioural tendencies associated with specific life history stages or dominance positions, which appear to be individual phenotypes because the data collection protocol does not extend across the lifespan of the species (34–36). Indeed, within the human literature, longitudinal studies suggest human behavioural tendencies and personality may be more labile than previously thought, with shifts in these tendencies predicted by a combination of age-related change or adjustment to particular life events, e.g. marriage (37–39). Distinguishing individual differences that are independent of life history or artefacts of socio-demographic variables requires long-term data reflecting changing life history within a particular study species.

The majority of non-human animal (hereafter animal) studies examining the repeatability of social behaviour have used comparatively short-term datasets, representing a limited period within the lifespan of the study species (34). In our study, we examine how social behaviours vary between individuals in wild, adult chimpanzees, utilising a behavioural dataset spanning over 20 years and more than 7500 full-day focal follows. We examine whether social behaviours (grooming, aggression and association) are repeatable over time both on a daily scale and when aggregated on a yearly level, with each sex analysed separately. By controlling for demographic and intrinsic factors, such as group size and age, we aim to determine the amount of variation in these social behaviours that is most likely attributable to individually stable phenotypes, thus limiting pseudo-repeatability (29).

Chimpanzees are an interesting study species to discern whether and how consistent individual differences in sociality arise and are maintained. These primates live in multi-female, multi-male groups with a high degree of fission-fusion dynamics (16,40), allowing potential for considerable fluctuation in social organisation during the course of their long lifespans (41). Chimpanzee societies also feature various cooperative behaviours, such as alliance formation, food sharing, group hunting, and territorial patrols, which likely impact on fitness and for which strong social relationships are required (42–45). Therefore, these individuals face diverse social environments and important choices regarding their social behaviour (46), which may predict a degree of flexibility in social tendencies.

In terms of dominance structure, when compared to female hierarchies, male hierarchies are dynamic and defined by high male-male competition, and there tends to be considerable reproductive skew towards high-ranking males (47–49). Although female dominance ranks do change during their lifespan, they are comparatively stable (50–53). Therefore, we expected a high degree of within-individual variation in aggressive tendencies in males in relation to lifetime fluctuations in dominance rank, whereas within-individual variation in aggression would be lower in females due to relatively stable hierarchy structure. Individual chimpanzees are highly strategic and flexible in grooming partner choice (46), while time devoted to grooming in this species can vary depending on demographic factors such as group size (54). Therefore, we expected low repeatability in grooming behaviour across long time periods. Lastly, chimpanzees sociality is characterised by a high degree of fission-fusion, allowing individuals to adjust to variation in within-group competition arising from ecological constraints, such as the availability of receptive mating partners or food (40). Competition will vary both seasonally and in the longer-term due changing group sizes or sex ratios. Therefore, we anticipated low repeatability for association behaviour as chimpanzees adjust to these fluctuations in competition.

## Methods

### Study Groups and Data Collection

Daily focal follow (55) data have been systematically collected by the Taï Chimpanzee Project, Côte d’Ivoire, since 1992 (56,57). We focussed on data collected since 1996, when data collection was consistent for behaviours relevant to this study and the control factors included in the models (see Statistical Analyses). This data includes observations of adult (>12 years) males and females from three fully habituated communities of chimpanzees: North (1996 – 2016), South (2002 – 2016), and East (2012 – 2016).

We collated these data to identify repeatability in interaction rates on two levels: daily and yearly (with year ending on 31^st^ August), allowing us to control for socioecologial variables at different temporal scales. For each of these levels, we restricted the dataset to individuals with regular focal follows to ensure that the data were sufficient to capture their typical social behaviour.

For the analysis on the daily level, we included focal follow days of adult individuals that lasted at least three hours, and included individuals for whom at least 10 focal follow days were available, resulting in a dataset of 70 individuals (45 females, 25 males) and 7615 individual focal follow days (individual mean = 109 days, max = 413 days).

For the analysis on the yearly level, we included adult individuals who were followed at least nine times in the time period from 1^st^ September to 31^st^ August and were present in their group for all this period, and who fulfilled these criteria at least in 3 years. This resulted in a dataset of 45 individuals (24 females, 21 males) and 272 years (individual mean = 6 years, max = 15 years). The number of years or days per subject of the study are detailed in Table S1 of the supplementary materials.

### Social Phenotype

On both the daily and the yearly level, we focused on three social variables: grooming, aggression, and association. We calculated variables from the perspective of the focal individual and with the focal as the actor (in case of the interactions) to ensure that they represent the individual’s social propensity as much as possible.

For grooming, we extracted the time (in minutes) focal individuals spent grooming adult partners (we focused on grooming given to others rather than overall time grooming, i.e. including grooming received, as this would reflect a tendency to attract grooming partners rather than an individual tendency to groom). For aggression, we extracted the number of aggressive interactions in which the focal individual was the aggressor. Both these variables were calculated the same for the daily and yearly level. We calculated association differently for the two levels: on the daily level, we extracted the cumulative number of individuals with whom the focal individual was seen in the same ‘party’ (defined as all individuals within visual contact of the observer – usually < 35 m) during that day. On the yearly level, as cumulatively individuals will likely associate with all other partners during this timeframe limiting variation, we instead calculated the total strength of connections of individuals by summing their association with all adult group members, using the simple ratio index (58) for that year based on common party membership of individuals. We standardized association strength by dividing it by group size.

As association indices have to be standardized by an appropriate null model (59), we conducted permutation analyses to confirm that associations were different than would be expected by random. For this analysis, we generated 1000 permutations of randomizations of party membership in which the number of individuals per party was kept constant and subsequent parties that originally had the same party membership also remained the same after randomization to account for autocorrelation (60). The overall number of times individuals appeared was not held constant, as we were interested in whether they would be more central/gregarious than expected. We subtracted the mean strength of individual’s connections with all other group members arising from the permutations from the observed strength, effectively creating an index of gregariousness (above 0 = more connected than would be expected in that group in that year; below 0 = less connected than expected).

### Statistical Analyses

All data were prepared and analyses were conducted using R 3.6.1 statistical software (61), with Generalized Linear Mixed Models (GLMM) fitted using the ‘lme4’ package (62). Models were fitted separately for males and females, for the dependent variables of aggression, grooming, and association, and for the daily and yearly levels of data aggregation, resulting in 12 models.

The **age** of all individuals was either known (for individuals who were not yet adult at the beginning of habituation) or was estimated in the beginning of the habituation period by experienced observers. Age ranged from 12 years (cut-off value) to 52 years, and the value assigned was either for the day (daily level) or the beginning of the yearly period (yearly level). We included age as a squared term in all models to account for potentially non-linear developmental patterns, e.g. individuals in their prime being more aggressive than younger and older individuals.

**Dominance rank** was calculated using a modification of the Elo rating method (63,64) based on unidirectional pant grunt vocalizations within each sex and standardized between 0 and 1 within each group. For the daily level, rank was extracted for each day of data an individual was observed. Rank was extracted on 31^st^ August for each yearly level.

For females, as **reproductive state** and **infant age** influences gregariousness and sociality (65–69), on the daily level, we included a factor with five levels: maximum tumescence of sexual swellings, mother with a new-born infant (below 3 months of age), mother with an un-weaned offspring (below 4 years of age), mother with weaned but immature offspring, or none of the above. In instances where females had weaned or un-weaned offspring but were also maximally tumescent or had a new-born infant, they were classified as maximally tumescent or with a new-born infant, as these reproductive states were anticipated to have a greater effect on behaviour than the presence of older offspring (81–85). On the yearly level, we included whether they had no offspring throughout the year, had a new-born at any point, or un-weaned offspring. On the daily level for the analyses concerning males, we included the *number of females with maximally tumescent sexual swellings* on that day as a variable.

As the **sex ratio** of adult individuals in a group can influence social behaviour, we calculated the adult sex ratio (number of males/number of females) of each group at the start of the yearly period (for the yearly level) or of those individuals present in the focal individual’s party during the day (for the daily level).

As interaction rates could be determined by the **number of available partners**(on the daily level) or **group size**(on the yearly level), we included these variables into the models. The yearly group size variable was standardized within groups, as there would otherwise be complete separation between the group identity and group size (as North group was considerably smaller than the others).

For both daily and yearly levels of analysis, we included an offset term for focal observation time (in hours; log-transformed) to account for differences in base rates of how often individuals could be observed interacting. We accounted for seasonality effects in the daily level analyses by including the radians of Julian date as a control variable (70). All models included a random effect of group by year: as group dynamics change over time, and we do not expect the three communities to show the same interaction rates, this term accounts for overall differences that could be influencing how social individuals are.

### Individual Model Structures

For **aggression**(both on the daily and yearly level, and males and females), we fitted GLMMs with Poisson error structure and log link (71), using the number of bouts per day/year as outcome variable, and having a log-transformed offset term for observation time to control for differences in observation effort (72). For the daily level, we included age as a quadratic term, dominance rank, available partners (daily level) or group size (yearly level), sex ratio, and group identity as fixed effect predictors. For the daily level, we also included the number of females with maximum swellings and the cosine and sine functions to capture seasonal effects. We included the random intercept of Year within Group, to account for the fact that data collected in the same time period in the same group are not independent, and the random intercept of Individual Identity. We also included the random slopes for rank, age, available partners, and sex ratio in Individual Identity, and rank and available partners in Group Year (73).

For **grooming**, on both the daily and yearly level, we fitted LMMs with Gaussian error structure, using the log-transformed hourly grooming rate as outcome variable. The fixed effects and random effect structure were the same as described for the aggression models.

For **association** on the daily level, we fitted GLMMs with Poisson error structure, using the number of individuals that the focal associated with on the day as outcome variable, and the number of available partners in the group as offset term. The fixed effects were the same as described for aggression, bar the removal of the available partners from the fixed effects. For gregariousness on the yearly level, we used the difference between observed and expected association strength as outcome variable, fitting LMMs with Gaussian error structure, and the same fixed effects as described above.

For all models, we z-standardized continuous predictor variables to facilitate interpretation (74). We compared the full models including all predictors and random intercepts and slopes with a null model including all fixed and random effects except the random intercept and slope for the individual identity (19,75), to test whether stable individual differences had an impact on interaction distributions. To establish the strength of the observed effect of individual identity only (76), we applied the ‘r.squaredGLMM’ function of the ‘MuMIN’ package (77) on a reduced model containing only the fixed effects and the random intercept and slopes for individual identity. We assumed that the variance in social behaviour that can be attributed to intra-individual stability can be captured by subtracting the variance explained by the fixed effects alone (marginal effect size) from that explained by the fixed effects and the individual random effect structure (conditional effect size) (76,78). Thus, for both the daily and the yearly level, both sexes, and all three social behaviours, we report whether the random effect significantly improved predictability, the marginal and conditional effect size, and their difference.

To avoid problems due to multicollinearity, we established the Variance Inflation Factors (VIF) of each model (79) using a simple linear regression of the fixed effects and applying the ‘vif’ function of the ‘car’ package (80). The maximum VIF for any model was 3.13, indicating that collinearity was not an issue. For all Poisson models, we tested for overdispersion, which was not an issue in any model.

## Results

We report the effect sizes attributed to the individual random effects and the comparison of the full model containing the random effect and the null model without it for the different models in Table 2 and Figure 1.

**Table 2:** Repeatability coefficients of chimpanzee social behaviours. Effect sizes and results of full null model comparisons for the models testing the impact of the individual random intercept and slopes on grooming rates, aggression bouts, and association in male and female Taï chimpanzees on a daily and yearly level. The correlation between the two levels was conducted on the random intercept residuals of individuals who were present in both datasets.

|  |  |  | Marginal R<sup>2</sup> | Conditional R<sup>2</sup> | Individual R<sup>2</sup> | X<sup>2</sup><sub>df = 5</sub> | p | Correlation<br>Daily – Yearly |
| --- | --- | --- | --- | --- | --- | --- | --- | --- |
| Female | grooming | daily | 0.08 | 0.26 | 0.19 | 50.87 | <0.001 | 0.73 |
|  |  | Yearly | 0.20 | 0.60 | 0.40 | 44.42 | <0.001 |  |
|  | aggression | Daily | 0.11 | 0.25 | 0.14 | 13.55 | 0.019 | 0.70 |
|  |  | yearly | 0.17 | 0.67 | 0.50 | 13.04 | 0.011 |  |
|  | association | daily | 0.25 | 0.67 | 0.42 | 450.93 | <0.001 | 0.30 |
|  |  | yearly | 0.11 | 0.72 | 0.61 | 36.29 | <0.001 |  |
| Male | grooming | daily | 0.10 | 0.28 | 0.18 | 50.71 | <0.001 | 0.93 |
|  |  | yearly | 0.33 | 0.74 | 0.42 | 28.32 | <0.001 |  |
|  | aggression | daily | 0.30 | 0.54 | 0.24 | 150.69 | <0.001 | 0.55 |
|  |  | yearly | 0.67 | 0.93 | 0.26 | 202.19 | <0.001 |  |
|  | association | daily | 0.47 | 0.83 | 0.35 | 71.08 | <0.001 | 0.20 |
|  |  | yearly | 0.24 | 0.45 | 0.21 | 9.61 | 0.022 |  |

**Figure 1:**
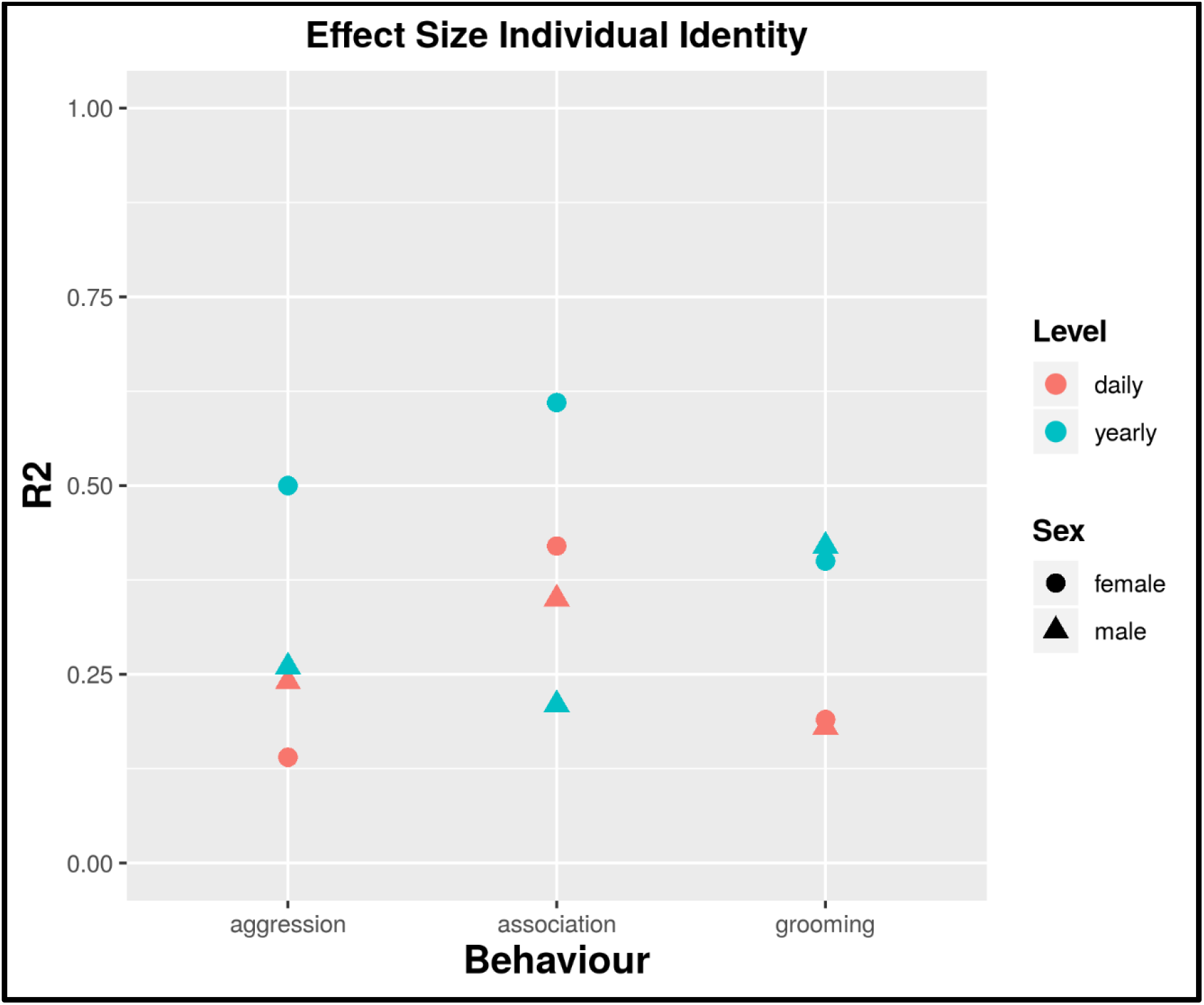
Repeatability of Chimpanzee Social Behaviours. Overview of the effect sizes attributed to the individual random effect, delineated by behaviour, sex, and timeframe (yearly vs daily) over 20 years of data.

In all models, including the random intercept and slopes of individual identity significantly improved model fit, indicating that for all social behaviours involved in this study, chimpanzees showed inter-individual differences that were consistent over time and could be detected on the daily and yearly level.

The variation explained by the random effect of individual identity varied between 0.14 and 0.61, and inter-individual differences tended to be more pronounced on the yearly level (mean = 0.40) than on the daily level (mean = 0.25). Thus, knowing the individual identity, for example, would allow one to predict 50% of variation in female aggression rates across years (Table 1). For grooming, the results for both sexes were almost identical in terms of the impact of the random effect of identity (daily = 0.18/0.19, yearly = 0.40/0.42). In general, inter-individual differences between females were more stable than between males (overall female mean = 0.38 vs overall male mean = 0.28), especially on the yearly level (female yearly mean = 0.50 vs male yearly mean = 0.30). Thus, after accounting for age, dominance rank, reproductive state (for females), group-level sex ratio and group size, stable inter-individual differences in yearly interaction rates where more pronounced in females than in males. For grooming and aggression, the residuals of individuals that were represented in both the yearly and daily models were highly correlated (see Table 1), while this was not the case for gregariousness, where different outcome variables were used. Figures 2 and 3 illustrate inter-individual variation in grooming, aggression and association at the daily and yearly levels in male and female chimpanzees respectively.

**Figure 2:**
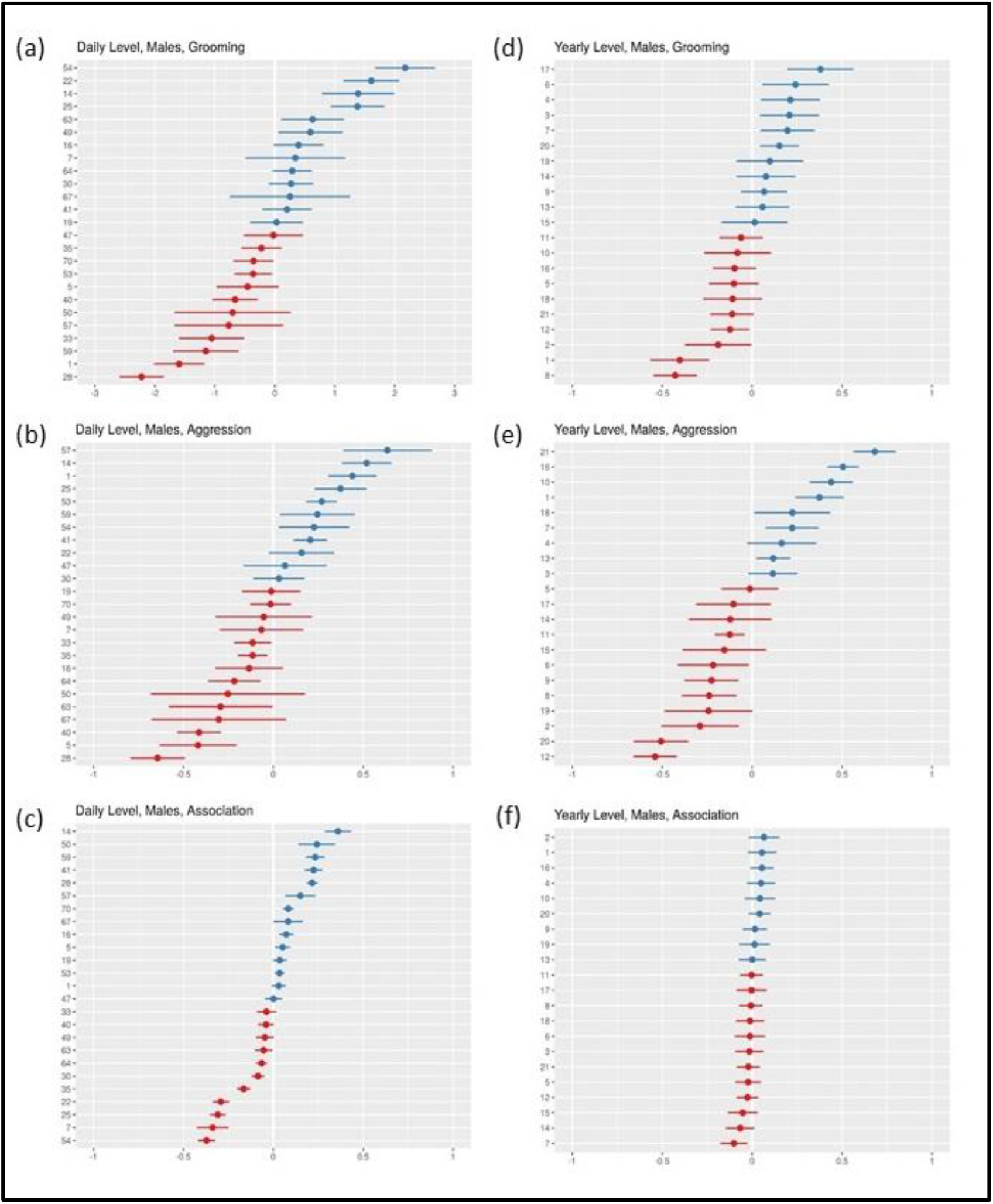
Inter-individual variation in grooming, aggression and association in male chimpanzees. Plots include random effect coefficients for individual identity from models examining variation in grooming, aggression and association in male at both the daily (a, b, c) and yearly (d, e, f) level of aggregation. The x-axis indicates the random effect coefficient; the y-axis indicates individual identities. Blue individuals expressed the variable of interest (grooming, aggression, and association) more than the average expression of the variable given all fixed and random effects within the model, red individuals expressed the variables less than the given average.

**Figure 2:**
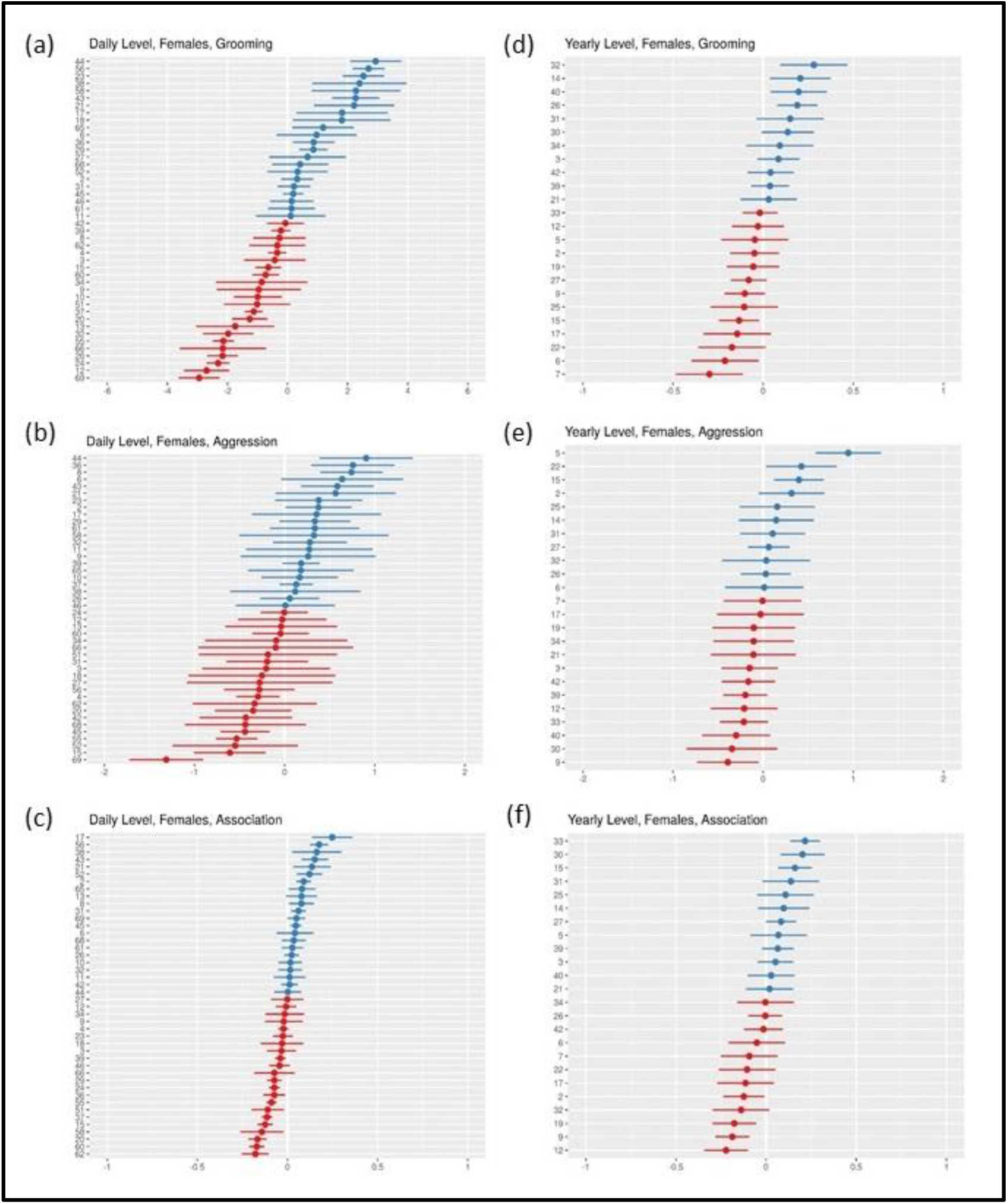
Inter-individual variation in grooming, aggression and association in female chimpanzees. Plots include random effect coefficients for individual identity from models examining variation in grooming, aggression and association in male at both the daily (a, b, c) and yearly (d, e, f) level of aggregation. The x-axis indicates the random effect coefficient; the y-axis indicates individual identities. Blue individuals expressed the variable of interest (grooming, aggression, and association) more than the average expression of the variable given all fixed and random effects within the model, red individuals expressed the variables less than the given average.

## Discussion

Our study reveals a high degree of repeatability in social behaviour in wild chimpanzees over several years and in many of our subjects, over a sizeable proportion of the adult lifespan of the species. The repeatability estimates for social behaviour in our study were comparable to an average repeatability estimate of behavioural traits across numerous animal taxa formerly generated by a meta-analytical study (R^2^=0.37; (81)). Importantly, unlike many former studies, using a comprehensive dataset, we were able to control for temporal, seasonal and demographic changes across the long-term study period that may influence variation in social behaviours, and thus limit pseudo-repeatability (29). As such, our method provides reasonable confidence that our repeatability estimates reflect stable social behaviour phenotypes that may be independent of life history stage or strategy.

The three behaviours of aggression, grooming (affiliation) and association represent three important components of sociality for chimpanzees, facilitating social goals such as dominance rank attainment (82,83) and the formation of social bonds (42,84), both of which influence fitness (43,44,85–88). Our study expands on former research in this field in terms of its scale, both temporally and in the range of behaviours examined, in a wild population, demonstrating that social phenotypes are a phenomenon in long-lived species occupying complex social environments. Indeed, given the results of our study and work conducted in taxa ranging from insects (23) to fish (24,25) to birds (19) to primates (20,89), consistent individual differences in social phenotypes seem the norm for group-living animals regardless of the degree of social complexity.

Repeatability was generally lower in the daily versus the yearly level, with implications for data aggregation and interpretation in repeatability analyses. The daily measures incorporate more data points with potentially fewer social interactions within them compared to the yearly measures. This may result in more random error and thus lower repeatability estimates for the daily level. Nevertheless, the differences between the daily and yearly measures are still informative.

The contrast between daily and yearly levels of repeatability was strongest in female aggression, with much higher repeatability estimates in the latter. Male aggression repeatability was low for both the daily and yearly levels. This likely reflects sex-differences in dominance hierarchies. Males use aggression to improve their position within the dominance hierarchy and to increase their reproductive opportunities (90–93). Therefore, their aggression rates can fluctuate due to variation in hierarchy stability and male-male competition (14,92). Compared to males, female chimpanzee hierarchies are relatively stable (51,53) and female aggression rates are less likely to be determined by variability in these factors. Instead, female aggression in chimpanzees is more likely determined by individual tendencies and seasonal, but comparatively predictable, variation in food availability (51). However, aggression is a costly behaviour in terms of energetics (94,95) and comes with inherent risks of injury, which could even be lethal in certain chimpanzee populations (96,97). Therefore, although in the long-term certain females are more aggressive than others, on a day-to-day basis, individuals are likely to be selective in when to be aggressive, reducing inter-individual differences and repeatability measures.

Females had higher repeatability estimates than those calculated for males for association. However, for both sexes, within-individual variation appeared much lower for association compared to the other social measures. We did not observe sex differences in the repeatability estimates for grooming and within-individual variation in this behaviour again seems comparable for each sex. Compared to other populations, Taï chimpanzees are considered highly gregarious, with the contrast in this population-level difference most apparent when examining female gregariousness (65). Our results suggest that these chimpanzees have relatively invariable preferential party sizes and rates of association. While grooming forms an important component of social bond formation (84), chimpanzees also make contingent grooming choices based on a range of parameters, such as audience, partner rank or context (e.g. reconciliation after an aggression) (68). Therefore, our results highlight this contrast in social decision making between levels of association (constrained preferences) and grooming (flexible choices).

Importantly, we found consistent individual differences in multiple forms of social interaction. Much research to date on social phenotypes has focused solely on association patterns (19,25), or on single forms of dyadic interaction (20, 34), with a minority of studies examining consistency in multiple forms of social behaviours (89,98,99). Our study shows that consistent individual differences in social behaviour extends to patterns of aggression and affiliation, both of which should influence fitness more than association alone (43,44,85–88,100). Indeed, both aggression and grooming involve direct, typically physical interactions with other group members, meaning that variation in these phenotypes will be important for factors such as rank acquisition (51,64,68,101,102), disease transmission (103–105) and group cohesion (45,106,107). Given the potential significance of these social tendencies, understanding how certain individuals come to be more aggressive or affiliative, as well as gregarious, than others, requires further empirical exploration.

The social niche hypothesis proposes that individuals adopt particular social strategies to ameliorate the resource competition based on individual idiosyncrasies that might predict competitive ability, such as body size (33). Chimpanzees have a protracted immature period, lasting around 10-15 years, during which, young chimpanzees consistently associate with their mothers (108). That some adult females (which will also typically be mothers) are more aggressive, affiliative and gregarious than others is significant to offspring development. Given repeatability, mothers consistently expose their offspring to particular environments, whether more or less social (109,110). In certain chimpanzee populations, high-ranking females tend to occupy areas within a group’s territory with more food resources, with low-ranking females typically occupying more resource poor areas (50). Furthermore, high-ranking chimpanzee mothers tend to have larger offspring (111), a pattern evident in our study population (112). This suggests that young chimpanzees may have to adopt particular social strategies arising from both their experience of competition via social exposure and their own competitive ability, which may also be determined by maternal effects. Therefore, having revealed that stable social phenotypes can be maintained over years in one of our closest living relatives, chimpanzees are promising model species to disentangle the genetic and environmental contributions to these phenotypes by taking a developmental approach to their emergence in younger subjects.

Our approach tested for stable individual differences in social tendencies that are independent of socioecological influences on the rates of behaviour. As a growing number of longitudinal studies have highlighted, age- or life event-related changes in behavioural tendencies are apparent in humans (37–39). One approach to further characterise and understand social phenotypes in chimpanzees may be to explore whether certain social phenotypes respond to comparable life events that occur within the species and to what degree. For example, the stable social phenotypes generated in the present study may predict behavioural and physiological responses to the loss of positions within a dominance hierarchy. This would allow the testing of consistent individual differences in *plasticity* to changing environments or internal state and provide insight on the mechanisms maintaining individual differences in social phenotypes over time.

In summary, our results add to the well-established literature on the repeatability of behaviour and social tendencies in group-living animals. Our study informs specifically within chimpanzees on important sex-differences in social tendencies. Furthermore, given the long development period in this species, our results present exciting opportunities to explore in detail the factors that contribute to the emergence of social phenotypes, such as via social niche specialisation.

## Acknowledgments

We thank the Ministère de l’Enseignement supérieur et de la Recherche scientifique and the Ministère des Eaux et Forêts of Côte d’Ivoire, and the Office Ivoirien des Parcs et Réserves for permission to conduct the study. We are grateful to the staff of the Taï Chimpanzee Project and the Centre Suisse de Recherches Scientifiques for support. We are indebted to the numerous field staff that have collected the data presented.

